# Benchmarking MSA pairing for protein-protein complex structure prediction reveals a depth-over-pairing principle

**DOI:** 10.64898/2026.04.14.718427

**Authors:** Yuxian Luo, Wenkai Wang, Zhenling Peng, Jianyi Yang

## Abstract

AlphaFold-Multimer (AFM) and AlphaFold3 (AF3) have revolutionized protein-protein complex structure prediction. However, it remains unclear whether elaborate MSA pairing is a strict prerequisite for achieving high accuracy. Here, we systematically evaluated diverse MSA construction strategies using a rigorous benchmark of 439 heterodimers. We found that incorporating paired MSAs yields only marginal improvements in structure prediction. Crucially, experiments using shuffled pairing patterns reveal that these gains stem not from specific pairing constraints, but rather from the increased depth of input MSAs, a finding that also extends to higher-order multimers. Furthermore, we demonstrate that prioritizing the inclusion of more homologs without pairing proves to be a superior strategy, enhancing performance for both intra- and inter-species dimers as well as the more challenging antibody-antigen complexes. This observation is substantiated by the recent success of the omicMSA strategy. Mechanistic analysis indicates that this capability is driven by inter-subunit physicochemical complementarity and the intrinsic architecture of the AF3 network. Furthermore, we identify critical bottlenecks that limit prediction accuracy, including large complex dimensions, restricted interface areas, and low experimental resolution. These findings establish a “depth-over-pairing” principle, offering new insights for improving protein complex prediction.

## INTRODUCTION

Protein-protein interactions (PPIs) govern the vast majority of cellular processes, making the elucidation of their quaternary structures critical for both mechanistic biology and drug discovery [1]. In recent years, this field has been revolutionized by deep learning-based prediction methods, with algorithms such as AlphaFold-Multimer (AFM) [2], RoseTTAFold2 (RF2) [3], and AlphaFold3 (AF3) [4] achieving unprecedented accuracy. Central to the success of these models is their ability to effectively leverage evolutionary signals derived from Multiple Sequence Alignments (MSAs). Leveraging the principle that interfacial residues coevolve to preserve interaction compatibility [5], coevolutionary analysis has been integrated with structure prediction frameworks to enable proteome-scale identification of PPIs [6, 7]. This progress has been accelerated by the state-of-the-art lightweight deep learning networks designed for rapid PPI screening, which have achieved high accuracy across both bacterial pathogens [8] and the human proteome [9].

Traditionally, a prevailing paradigm in protein complex prediction is the construction of “paired” MSAs to capture inter-chain co-evolutionary information. Early methods for prokaryotic targets capitalized on genomic proximity (e.g., operon organization) to infer interactions [10]. For eukaryotic systems lacking such genomic context, the standard phylogeny-based pairing strategy (utilized by AFM and AF3) relies on concatenating homologs via strict species-level matching [11], often prioritizing candidates through sequence identity ranking. To augment the sensitivity and accuracy of these alignments, a diversity of advanced pairing strategies has emerged. For instance, approaches utilizing multiple taxonomic ranks have been proposed to relax species constraints, thereby recruiting remote homologs that are otherwise discarded [12]. With the advent of protein language models (PLMs), the field has seen further innovation: ESMPair refines phylogeny-based pairing by leveraging PLM embeddings to optimize the ranking of paralogs within species boundaries [13], while DiffPALM exploits the masked language modeling capabilities of the MSA Transformer to perform differentiable pairing directly from sequence context [14]. Most recently, DeepSCFold, an attention-based deep learning framework, incorporates sequence-derived structure complementarity and interaction probabilities to guide MSA construction, reporting significant gains over standard protocols [15].

However, despite these innovations, a fundamental ambiguity remains regarding the driver of prediction accuracy in the era of AF3. Notably, as demonstrated by our participation in CASP15 [16] and CASP16 [17], our modeling strategy, which operates entirely independently of MSA pairing, achieved top-tier performance in protein complex prediction. Consistent with this observation, AF2Complex does not require paired MSAs and achieves a significant improvement over AFM [18]. It is thus uncertain whether the benefits of complex pairing algorithms are artifactual or essential. Currently, a systematic evaluation of these input dependencies is lacking. This gap forces the community to rely on suboptimal protocols for difficult targets characterized by a scarcity of paired homologs, e.g., antibody-antigen complexes.

To resolve this ambiguity, we conducted a systematic evaluation of the MSA pairing effect in AF3 using rigorous benchmarks. We demonstrate that AF3 is remarkably robust to the disruption of MSA pairing, with prediction accuracy driven primarily by MSA depth rather than explicit pairing. We further validate this observation through mechanistic analysis of inter-subunit complementarity and the network architecture, confirming that the model’s intrinsic capacity to extract latent co-evolutionary patterns bypasses the need for the paired MSA. Furthermore, we identify critical bottlenecks, such as large complex size, minimal interface area, and low experimental resolution, that continue to limit current performance. Our findings establish a ‘depth-over-pairing’ principle, offering new guidelines for efficient complex prediction and the training of next-generation models.

## RESULTS

### Does MSA pairing really contribute to complex structure prediction?

To rigorously quantify the contribution of MSA pairing to protein-protein complex structure prediction, we designed four complementary MSA input strategies (Fig. 1a). ***m*MSA** (‘monomer’): Standard monomeric MSAs for each subunit, concatenated in native sequence order without any inter-chain pairing. This serves as the unpaired baseline. ***p*MSA** (‘pair’): The *m*MSA supplemented with Species-paired MSAs. This matches the default protocol in AF3 and exhibited the strongest performance in our initial benchmarking of six pairing methods. ***s*MSA** (‘shuffle’): Serves as a control where the paired sequences from *p*MSA are randomly shuffled. This strategy preserves overall sequence composition, MSA depth, and monomeric co-evolutionary information while deliberately disrupting inter-chain co-evolutionary signals. ***u*MSA** (‘UniProt’): Comprises the *m*MSA merged with the full, raw UniProt MSAs (the source pool used for constructing *p*MSA). These sequences are concatenated without explicit pairing constraints. This tests whether simply increasing MSA depth and diversity—without explicit pairing—can recapitulate the benefits of deliberate pairing in *p*MSA. Full details of MSA sources, construction pipelines, and hyperparameters are provided in the Methods section.

**Fig. 1.**
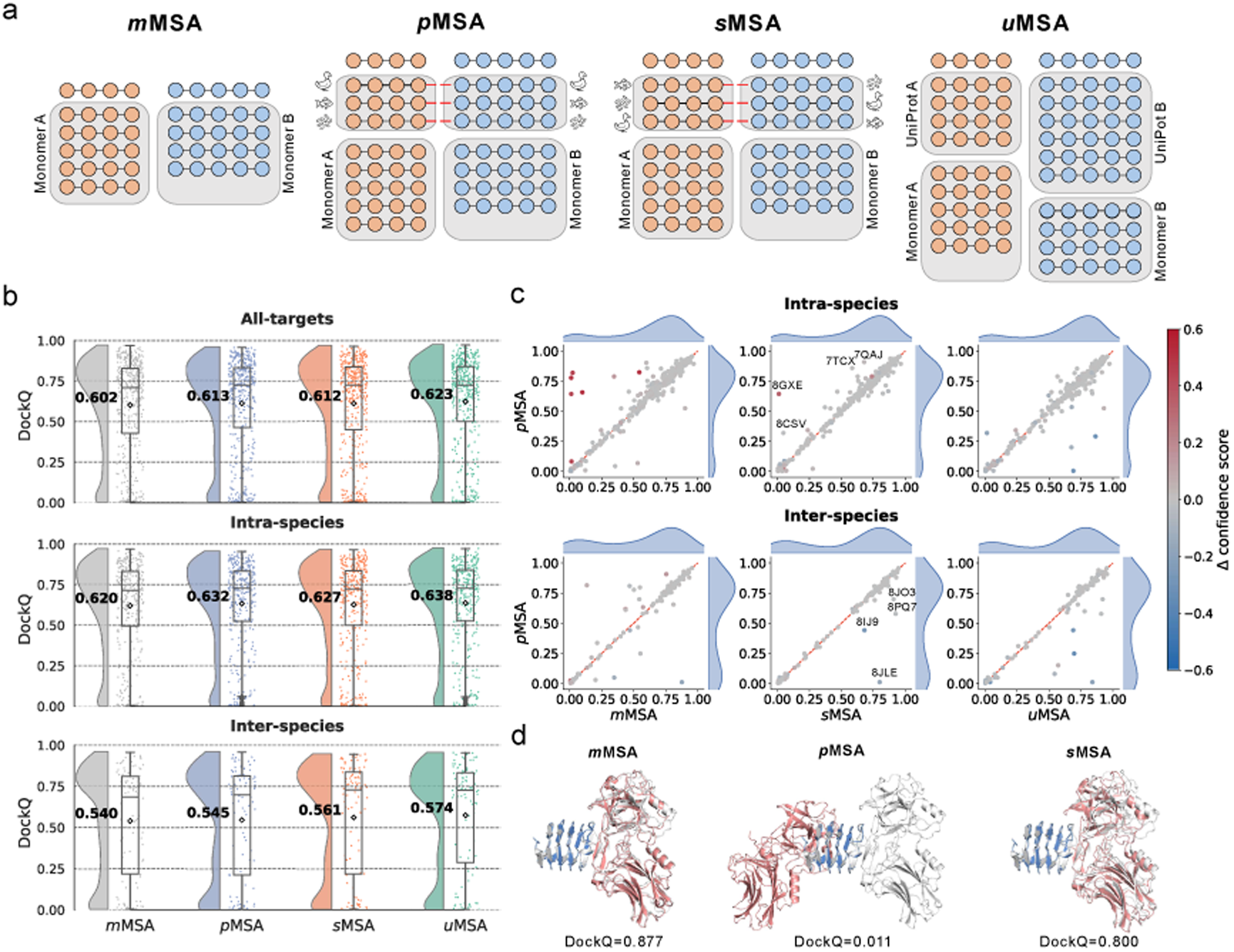
Schematic of MSA construction strategies and performance on the HD439 dataset. (**a**) Schematic illustration of the four MSA input strategies: *m*MSA, *p*MSA, *s*MSA, and *u*MSA. Red dashed lines denote sequence pairing (shown for Species-based pairing as an example). (**b**) Distribution of DockQ scores for *m*MSA, *p*MSA, *s*MSA, and *u*MSA strategies, evaluated on the full HD439 dataset as well as on intra-species and inter-species subsets. Diamonds and bold numerals indicate mean DockQ scores. (**c**) Head-to-head scatter plots comparing *p*MSA (*y*-axis) against *m*MSA, *s*MSA, and *u*MSA (*x*-axis) for intra- and inter-species complexes. The color scale represents the difference in confidence score (calculated as *p*MSA minus the alternative method). (**d**) Structural superposition of top-ranked models predicted by AF3 using *m*MSA, *p*MSA, and *s*MSA inputs against the experimental structures for a representative target (PDB ID: 8JLE). Experimental structures are shown in gray cartoon, while predicted structures are colored by chain (A: blue, B: salmon).

We assembled a high-quality, non-redundant benchmark set of 439 heterodimeric protein complexes (**HD439**) that share no significant sequence similarity with the AF3 training set (see Methods for construction criteria). Given that the default pairing protocols in AFM and AF3 rely predominantly on species information, we further partitioned HD439 into 340 intra-species and 99 inter-species complexes to enable a detailed, context-specific analysis.

#### Inter-species complexes are more challenging to predict than intra-species complexes

Inter-species complexes proved substantially more challenging to predict than intra-species ones (Fig. 1b). For example, the *p*MSA yielded a mean DockQ of 0.545 for inter-species targets, significantly lower than the 0.632 observed for intra-species complexes (two-sided Wilcoxon rank-sum test, *P* = 6.76×10^−7^). This gap largely arises because inter-species interactions generally lack the strong, persistent co-evolutionary signals characteristic of intra-species complexes. Such signals are typically weaker or absent in transient or evolutionarily recent interfaces (e.g., host–pathogen or antibody–antigen interactions). Consistent with this, inter-species complexes exhibited markedly shallower paired MSAs, as quantified by the number of homologous sequence pairs (Supplementary Fig. 1a). The resulting sparsity of co-evolutionary information impairs the model’s ability to accurately infer inter-chain contacts and interfacial geometry, thereby exacerbating the prediction challenge for inter-species complexes.

#### Disrupting pairing relationships does not compromise AF3 modeling accuracy

Inclusion of the default paired MSAs (*p*MSA) produced only modest gains over the monomeric baseline (*m*MSA) across the HD439 dataset (mean DockQ: 0.613 vs. 0.602), with a slightly larger benefit for intra-species complexes (0.632 vs. 0.620; Fig. 1b). These results indicate that supplementing monomeric MSAs with paired homologous sequences contributes to improved prediction performance. That is consistent with previous reports in the literature and is the reason why substantial efforts were dedicated to optimizing MSA pairing algorithms, but with limited success.

Remarkably, however, our analysis reveals that this improvement does not stem from the preservation of explicit inter-chain pairing relationships. Randomly shuffling the paired sequences (*s*MSA; see Fig. 1a)—which disrupts co-evolutionary coupling signals while exactly preserving sequence composition, MSA depth, and monomeric co-evolution — yielded virtually identical overall performance to *p*MSA (mean DockQ: 0.612; *P* = 0.96; Fig. 1b).

For inter-species complexes, *s*MSA even outperformed *p*MSA (mean DockQ: 0.561 vs. 0.545). This counterintuitive result suggests that species-based taxonomic matching can introduce erroneous pairing constraints, particularly for chains with limited or disjoint evolutionary histories. Such mismatches generate spurious co-evolutionary noise that partially offsets the benefit of increased sequence depth. Head-to-head comparisons (Fig. 1c) show that *s*MSA maintains comparable or superior performance for most inter-species targets and yields substantial improvements in several cases (e.g., PDB entries 8JLE, 8IJ9, 8PQ7, and 8JO3; Supplementary Fig. 2b). A representative example is the inter-species complex 8JLE (Chain A: *Homo sapiens*; Chain B: *Clostridium botulinum*; Fig. 1d). While *m*MSA produced an accurate model (DockQ > 0.8), *p*MSA generated a grossly incorrect pose (DockQ ≈ 0.0), demonstrating that species-based pairing introduced misleading co-evolutionary signals. Shuffling (*s*MSA) fully rescued the prediction (DockQ = 0.800).

For intra-species complexes, *s*MSA showed a minor, non-significant decrease in mean DockQ relative to *p*MSA (0.627 vs. 0.632; *P* = 0.83; Fig. 1b). Closer inspection revealed that this small gap arises predominantly from inconsistencies in AF3’s built-in ranking score rather than from reduced modeling capability. In the four targets where *s*MSA underperformed *p*MSA by >0.2 in top-1 DockQ (8GXE, 8CSV, 7TCX, and 7QAJ; Supplementary Fig. 2a), high-accuracy models (comparable to *p*MSA) were nonetheless generated but ranked lower. These observations underscore the need for improved confidence estimation and model ranking methods in multimeric structure prediction [19].

Collectively, these findings demonstrate that explicit inter-chain pairing is not essential for enhancing AF3 performance in dimer structure prediction. Instead, the primary benefit of *p*MSA and *s*MSA derives from the expanded pool of homologous sequences. This is vividly illustrated by target 8DYE (Fig. 2a, d): *m*MSA failed to produce an accurate model (DockQ = 0.023) due to severely limited depth for Chain B (47 sequences), resulting in a distorted monomer (TM-score = 0.21) and incorrect quaternary arrangement. Supplementing with paired sequences (adding ∼1,450 sequences) increased Chain B depth to ∼1,500 regardless of pairing fidelity (*p*MSA or *s*MSA), restoring accurate tertiary structure (TM-score > 0.5) and enabling high-quality complex modeling (DockQ ≈ 0.8 in both cases; Fig. 2a). MSA depth, rather than pairing specificity, emerges as the dominant factor.

**Fig. 2.**
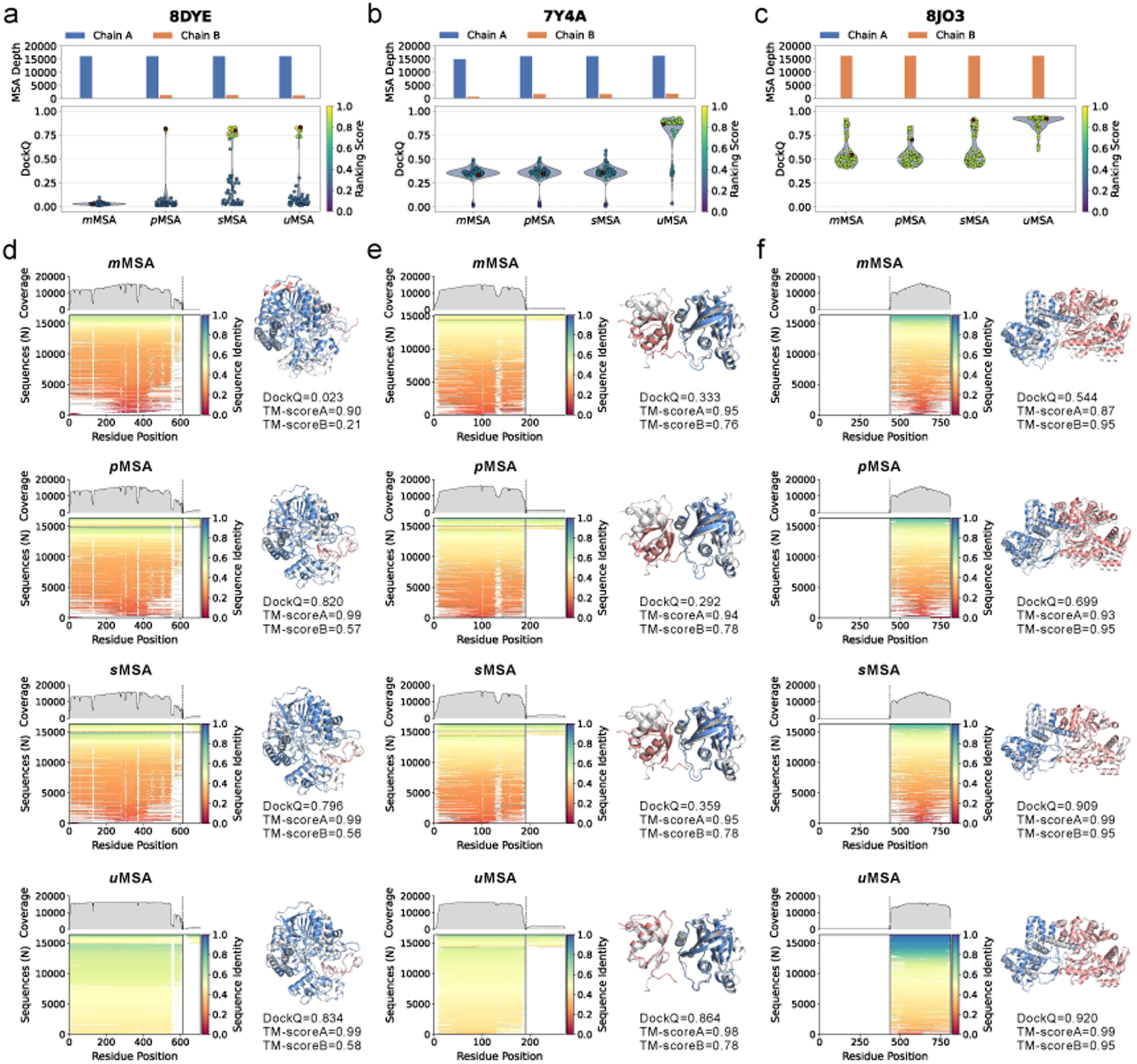
Representative case studies demonstrating the advantage of the *u*MSA strategy on complex structure prediction. (**a-c**) Quantitative analysis for targets 8DYE (a), 7Y4A (b), and 8JO3 (c). Top panels: Bar charts showing the effective MSA depth (number of sequences) for each chain. Bottom panels: Violin plots displaying the distribution of DockQ scores for 75 generated models. Red dots indicate the DockQ scores of the top-ranked models selected by the confidence metric. (**d-f**) Visualization of MSA characteristics and predicted structures for the corresponding targets in (a–c). Left panels: Heatmaps illustrating sequence identity (color gradient) and sequence coverage (top line profile) across residue positions. Right panels: Structural superposition of the top-ranked predictions (colored by chain; Chain A: blue, Chain B: salmon) onto the experimental reference (gray cartoon).

#### Inclusion of homologous sequences without pairing outperforms pairing methods

Motivated by the finding that MSA depth is more critical than pairing patterns, we sought to enhance performance by maximizing the inclusion of homologous sequences. To this end, we constructed *u*MSA (see Fig. 1a): a composite MSA merging the UniProt-derived monomeric MSAs (the raw sequence pool prior to pairing) with the standard *m*MSA, without any inter-chain pairing constraints. This approach retains the most extensive set of homologous sequences (Supplementary Fig. 1b), as it avoids the sequence loss often incurred during pairing procedures.

As anticipated, *u*MSA yielded the highest average DockQ score of 0.623 on the HD439 dataset, outperforming both *p*MSA and *s*MSA (0.613 and 0.612; Fig. 1b). Notably, this advantage was consistent across both intra- and inter-species complexes. To gain deeper insight into model behavior, we examined the full distribution of DockQ scores across all 75 generated models for the 20 examples exhibiting significant performance variations (ΔDockQ > 0.2) across the four MSA strategies (Supplementary Fig. 2). The corresponding MSA depths are displayed at the top of each plot. The results indicate that deeper MSAs in *u*MSA yield superior predictions. For instance, in targets 7THN, 7THQ, and 7M4M (Supplementary Fig. 2a), *u*MSA outperformed other methods in terms of both the top-ranked model and the overall DockQ distribution. The accompanying bar plots illustrate that *u*MSA provides substantially greater sequence depth for these three targets, underscoring the critical importance of MSA depth in modeling accuracy.

Beyond increasing absolute MSA depth, *u*MSA also enhances performance by improving MSA quality. For instance, for target 7Y4A (Fig. 2b, e), the *p*MSA already reached AF3’s practical sequence limit (16,384 sequences). However, *u*MSA—while constrained to the same input size—enriched the alignment for medium-identity homologs (sequence identity 0.4–0.6) and reduced low-identity noise (<0.2), dramatically improving prediction accuracy (DockQ from <0.4 to >0.8). A parallel improvement occurred for the target 8JO3 (Fig. 2c, f).

In summary, these results support the conclusion that enriching the pool of homologous sequences—rather than enforcing explicit pairing—is the more effective strategy for boosting protein complex structure prediction accuracy. This improvement stems from the enrichment of sequence information and the enhancement of MSA quality, which empower the network to sample high-accuracy models that were otherwise inaccessible due to the limited informational content of shallow, strictly paired alignments.

### Theoretical analysis for accurate complex prediction from unpaired MSAs in AF3

Although unpaired MSAs lack explicit inter-chain co-evolutionary signals, our results demonstrate that AF3 can nevertheless achieve high-accuracy predictions. We hypothesize that this is achieved through the synergy of two factors: physicochemical complementarity between subunits and the iterative updating mechanism in AF3.

#### Physicochemical complementarity between subunits

The extensive inclusion of unpaired sequences provides a rich evolutionary profile for each monomer, facilitating the accurate capture of the conserved surface patches, biophysical constraints, and intrinsic structural features. We hypothesize that AF3 leverages these monomer-specific signals to infer optimal docking poses based on geometric shape complementarity, electrostatic matching, and other physicochemical principles—thereby circumventing the need for explicit inter-chain co-evolutionary priors.

This mechanism is particularly evident in complexes with pronounced shape complementarity. A case in point is the *USP36–Ubiquitin-PA* complex (PDB ID: 8BS9; Fig. 3a, b), where the *USP36* catalytic domain presents a rigid, pre-formed binding cleft that recognizes *Ubiquitin* via a classic ‘lock-and-key’ fit. As a result, even the *m*MSA baseline, which lacks any inter-chain sequence pairing, yielded high-accuracy models (DockQ ≈ 0.9).

**Fig. 3.**
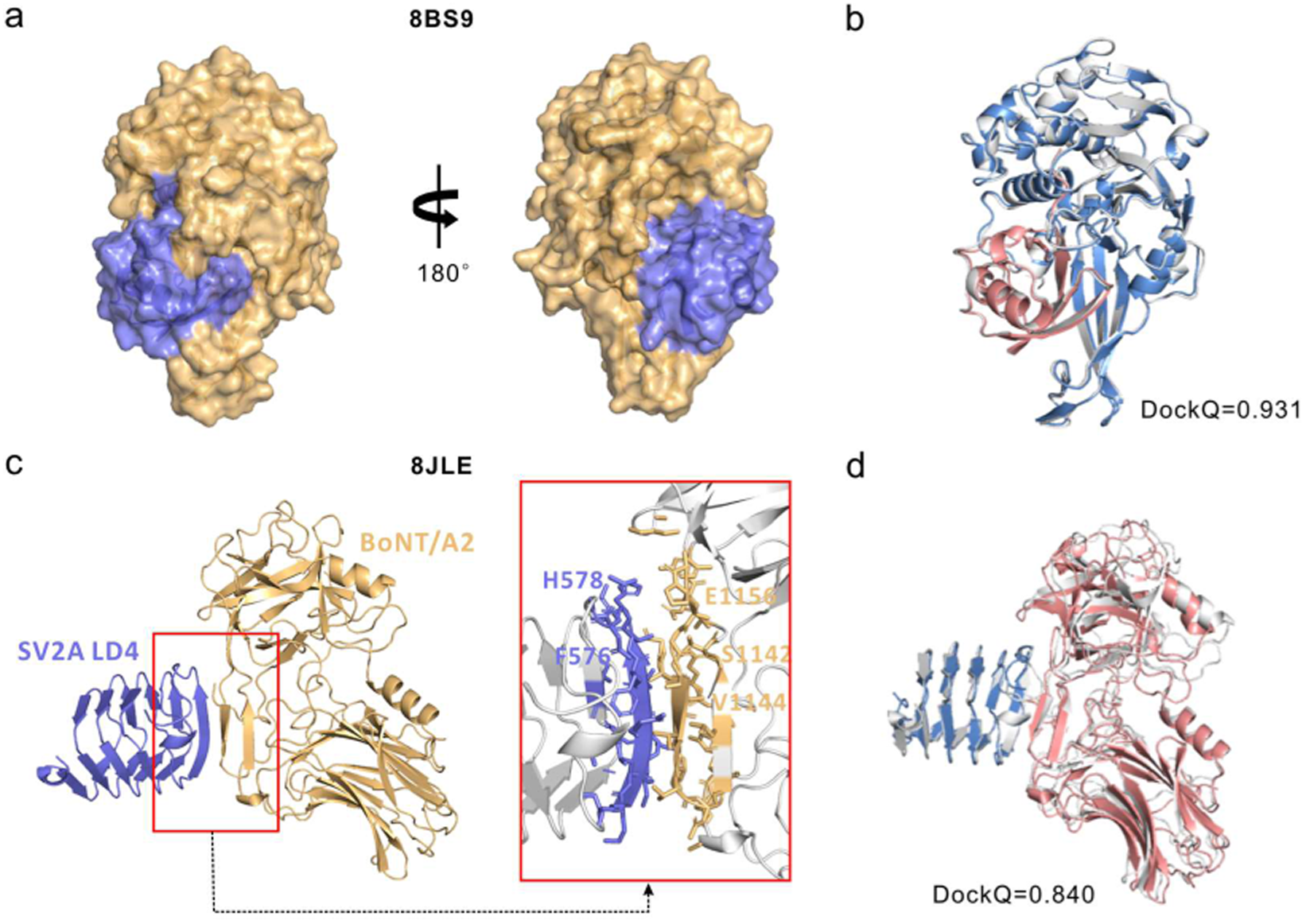
Examples of structural complementarity facilitating accurate predictions without explicit pairing. (**a**) Surface representation of the experimental structure for the USP36-Ubiquitin-PA complex (PDB ID: 8BS9), highlighting the shape complementarity between subunits. (**b**) Superposition of the AF3 structure predicted using *m*MSA (colored) and the native structure (gray) for 8BS9. (**c**) Experimental structure of the SV2A LD4 in complex with BoNT/A2 Hc (PDB ID: 8JLE). The inset provides a detailed view of the interaction interface with key residues labeled. (**d**) Superposition of the AF3 structure predicted using *u*MSA (colored) and the experimental structure (gray) for target 8JLE.

Another example is the *BoNT/A2–SV2A* complex (PDB: 8JLE; Fig. 3c, d). Structural analysis reveals that the interaction is mediated by the formation of a continuous intermolecular β-sheet (Fig. 3c), where the open edge of the BoNT/A2 β-hairpin ‘zips’ with the C-terminal β-strand of the SV2A LD4 β-helix via backbone hydrogen bonds [20]. This interface is reinforced by specific physicochemical complementarity, exemplified by a salt bridge between SV2A His578 and BoNT/A2 Glu1156, as well as the hydrophobic packing of SV2A Phe576 against BoNT/A2 Ser1142/Val1144 (highlighted in Fig. 3c). Leveraging these distinct structural features, AF3 successfully predicted the complex structure using the simple *m*MSA (DockQ = 0.877) and *u*MSA (DockQ = 0.840). Interestingly, the *p*MSA input resulted in a failed prediction (DockQ ≈ 0, Fig. 1d), aligning with our earlier observations regarding inter-species targets.

This case highlights the capacity of physicochemical compatibility to drive correct assembly independent of co-evolutionary signals.

Briefly, these results confirm that high-quality monomeric alignments can facilitate the accurate reconstruction of complexes through physicochemical complementarity, even in the complete absence of inter-chain co-evolutionary priors.

#### AF3’s iterative updating mechanism

Another critical factor lies in the network architecture of AF3. AF3 employs a conditional diffusion framework, which comprises two main components: the conditional component and the diffusion component. The former accounts for the majority of the computational load and plays a decisive role in structure prediction. Specifically, this component consists of the Template Module (2 blocks), the MSA Module (4 blocks), and the Pairformer (48 blocks). Given that templates were not utilized in our benchmark, our analysis focuses primarily on the MSA Module and Pairformer.

Specifically, MSA information is integrated and converted into the pair representation via the outer-product-mean operation within the MSA Module (Supplementary Eqs. (1) and (2)). Subsequently, this pair representation, encoding aggregated MSA signals, is updated through triangle multiplication and triangle attention. Crucially, both mechanisms leverage inner-product operations to facilitate dense interactions between any two positions within the MSA (Supplementary Eqs. (3) and (4)). In other words, this all-to-all interaction scheme enables the network to autonomously detect latent co-evolutionary patterns across the entire sequence space. Consequently, in principle, the model can reconstruct inter-chain evolutionary signals directly from raw unpaired alignments, bypassing the necessity for pre-defined pairing in the input MSA. A more detailed illustration of this mechanism is provided in Supplementary Note 1.

However, such initial, highly mixed signals require extensive refinement, a task performed by the deep Pairformer stack comprising 48 triangle update blocks. As shown in Supplementary Fig. 3, AF3 effectively captures inter-chain interactions primarily in the latter stages of the Pairformer. Interestingly, while the use of *p*MSA slightly accelerates this process for intra-species complexes (Supplementary Fig. 3a), the final performance converges to a level comparable to that of *s*MSA. For inter-species targets, the benefits of *p*MSA are negligible (Supplementary Fig. 3b). These results align with our theoretical understanding, confirming that the primary driver of performance is not the initial organization of the input MSA, but rather the network’s intrinsic capacity to extract latent interactions through deep iterative updating.

### The species-based MSA pairing outperforms other MSA pairing methods

To comprehensively understand the role of MSA pairing in protein complex structure, five additional pairing methods are benchmarked here. They were adapted from MULTICOM [21] and DeepSCFold [15] (see Methods for details). These include three annotation-based methods—pairing based on UniProt accession number proximity (**Distance**), known interactions from the STRING database (**STRING**), and co-occurrence in PDB entries (**PDB**)—and two protein language model (PLM)-based approaches that exploit either sequence similarity (**PLM**) or structural similarity (**PLMStr**). For each pairing method, the resulting paired MSA was combined with the standard monomeric MSAs of the individual subunits to generate input for AF3 structure prediction (similar to the construction of *p*MSA in Fig.1a, as described in “Monomer MSA generation”).

We compared the default species-based pairing against the five alternative strategies on pairwise common subsets of targets, as each alternative method is only applicable to a subset of complexes. This strict evaluation ensures a fair head-to-head comparison without confounding effects from differing target coverage. As shown in Fig. 4a, the default species-based pairing consistently achieved a small but consistent performance advantage across the evaluated alternatives, as measured by mean DockQ. These modest gains indicate that imposing additional pairing constraints—whether annotation-based or PLM-derived—does not substantially improve AF3’s prediction accuracy beyond the species-based baseline. Crucially, our key findings are robust across pairing methodologies. The superiority of *u*MSA (Fig. 4a) and the inherent modeling difficulty of inter-species complexes (Fig. 4b, c) persist regardless of the pairing strategy employed.

**Fig. 4.**
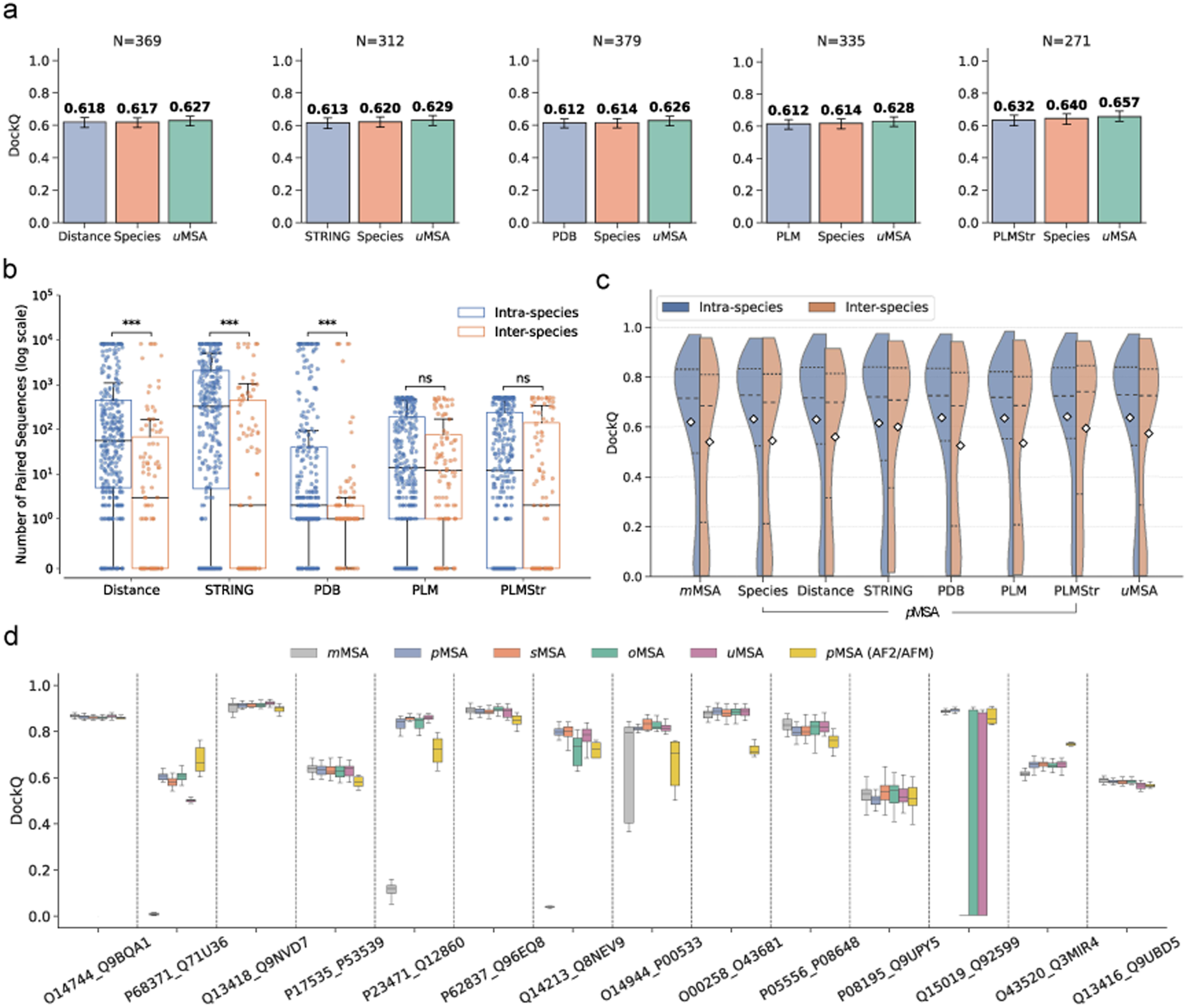
Analysis of alternative pairing strategies. (**a**) Performance comparison of five alternative pairing methods (Distance, STRING, PDB, PLM, and PLMStr) against the standard Species-based pairing and *u*MSA. *N* indicates the number of applicable targets within the HD439 dataset for each pairing method. Mean values are annotated above each bar. Error bars correspond to 95%. (**b**) Distribution of the number of paired sequences generated by alternative pairing methods for the intra- and inter-species subsets. (**c**) DockQ score distributions for AF3 predictions using alternative MSA pairing methods on the HD439 dataset, stratified by intra-species (blue) and inter-species (orange) complexes. White diamonds represent mean values. (**d**) Distribution of DockQ scores for 14 representative PPI targets for analyzing the effect of omicMSA. The evaluated MSA configurations in (d) are defined as follows: *m*MSA, comprising the residual unpaired sequences from the omicMSA pipeline; *p*MSA (i.e., the output of the omicMSA pipeline), consisting of species-based paired sequences supplemented with *m*MSA; *s*MSA (shuffled paired sequences supplemented with *m*MSA to break pairing relationships); *o*MSA (direct sequential concatenation of raw subunit MSAs from omicMSA, filtered at 95% identity to remove redundancy); and *u*MSA (standard alignments utilizing UniProt databases). *p*MSA (AF2/AFM) denotes the reference structural predictions (11 models per target) generated by AlphaFold2/AlphaFold-Multimer utilizing omicMSAs, as provided in [9].

### Evaluation of omicMSA in AF3 complex prediction

We further assessed the influence of the omicMSA strategy [9] on complex structure modeling. Compared to the alternative strategies that focus on pairing strategy, omicMSA is characterized by its extensive sequence database derived from unassembled genomic reads and draft eukaryotic genomes. The resulting MSAs offer sevenfold greater depth compared to those derived from UniRef100, thereby amplifying the detection of co-evolutionary signals for human PPI screening. We collected a dataset of 14 PPIs from the high-confidence predictions in [9], focusing on targets with experimentally determined dimeric structures deposited in the PDB (see Supplementary Table 1 for UniProt and PDB identifiers). Paired MSAs were generated from omicMSA using the official Colab notebook (https://colab.research.google.com/drive/1suholB5q6xn0APFHJE8cleMiCuv9gCk; “paired_unpaired” setting). Subsequently, three input configurations—*m*MSA, *p*MSA, and *s*MSA—were constructed based on the omicMSA results as outlined in Fig. 1a. Notably, the *p*MSA configuration represents the direct, unmodified output of the omicMSA pipeline. For each target, we generated 25 AF3 predictions across five random seeds. DockQ scores for the complete ensemble and the top-ranked models are provided in Fig. 4d and Supplementary Table 1, respectively.

Comparative analysis of average performance reveals that *p*MSA achieved a mean DockQ score of 0.756, significantly surpassing *m*MSA (0.611). Notably, for three specific targets (P68371_Q71U36, P23471_Q12860, and Q14213_Q8NEV9), the incorporation of paired sequences rescued predictions from incorrect (DockQ < 0.23) to medium- or high-quality predictions (DockQ > 0.50), accompanied by improved ranking confidence. Notably, the *p*MSA-driven AF3 predictions achieve greater accuracy than the AF2/AFM baselines established by the omicMSA authors (http://prodata.swmed.edu/humanPPI; DockQ: 0.756 vs. 0.739), confirming the general performance advantages of AF3 over AF2/AFM.

Crucially, *s*MSA achieved accuracy comparable to *p*MSA for most of the targets (13/14), indicating that the observed gains stem primarily from increased alignment depth rather than explicit pairing constraints, consistent with our earlier findings. Although the mean DockQ for *s*MSA (0.684) was lower than that of *p*MSA (0.756), this difference was largely driven by a single outlier (Q15019_Q92599; DockQ: 0.897 vs. 0.004), for which the baseline *m*MSA (without pairing information) had already achieved high accuracy (DockQ > 0.80). Moreover, the overall improvement of *s*MSA relative to *m*MSA further supports the conclusion that increasing sequence depth enhances complex modeling accuracy.

Note that *s*MSA can still imply ambiguous pairing relationships between the two sequence blocks. To bypass this potential pairing bias, we evaluated *o*MSA, which is constructed through a straightforward concatenation of the raw MSAs for each subunit generated by the omicMSA pipeline, completely free from any elaborate pairing constraints. Strikingly, *o*MSA achieved the highest mean DockQ score (0.764), marginally outperforming *p*MSA (0.756). This reinforces the conclusion that enhancing the quality of monomeric MSAs is a fundamentally more effective strategy than exploring complex pairing algorithms.

Additionally, we evaluated the previously defined *u*MSA for these 14 targets. The construction of *u*MSA strictly followed our standard methodology, relying on conventional databases (e.g., UniProt) instead of the expansive omicMSA sequence pools. While *u*MSA yielded robust results in the overall HD439 benchmark, it fell short of *o*MSA’s accuracy here (mean DockQ: 0.661 vs. 0.764). This disparity serves as direct evidence that the intrinsic quality of monomeric MSAs is the primary driver of prediction success.

In summary, the omicMSA evaluation highlights a clear trend: superior monomer alignments drive superior complex modeling. This strongly substantiates our core conclusion: improving monomer MSAs represents a fundamentally more impactful strategy than enforcing strict pairing constraints.

### Determinants of modeling failure in AF3

To identify the key factors limiting AF3’s accuracy for protein-protein complexes, we systematically analyzed the influence of MSA depth, geometric, biophysical, and experimental attributes on prediction performance across the HD439 benchmark.

#### MSA depth

Supplementary Fig. 4 illustrates the relationship between DockQ score distributions and MSA depth. As anticipated, increasing MSA depth elevates the performance ceiling, enabling AF3 to generate models with higher peak accuracy. However, a strict positive correlation is not observed across the entire distribution. Notably, while the deepest MSAs (> 10,000) are enriched for high-precision models, they also exhibited increased variance with a broader tail of low-scoring predictions. This suggests that while sufficient MSA depth is a prerequisite for high accuracy, the ultimate prediction success is likely governed by other intrinsic determinants beyond sequence abundance.

#### Protein length

Prediction accuracy declines markedly with increasing complex size (Fig. 5a). Complexes in which both subunits exceed 1,000 residues consistently yield low DockQ scores (< 0.1), irrespective of MSA strategy (*m*MSA, *p*MSA, *s*MSA, or *u*MSA). Representative failures include targets 8C8G, 8BYP, and 8PBY (Fig. 5d), where all models collapse to incorrect poses. This bottleneck likely stems from AF3’s crop size limit (maximum 768 tokens) during training, which restricts the model’s ability to capture long-range dependencies and intricate binding topologies in large assemblies.

**Fig. 5.**
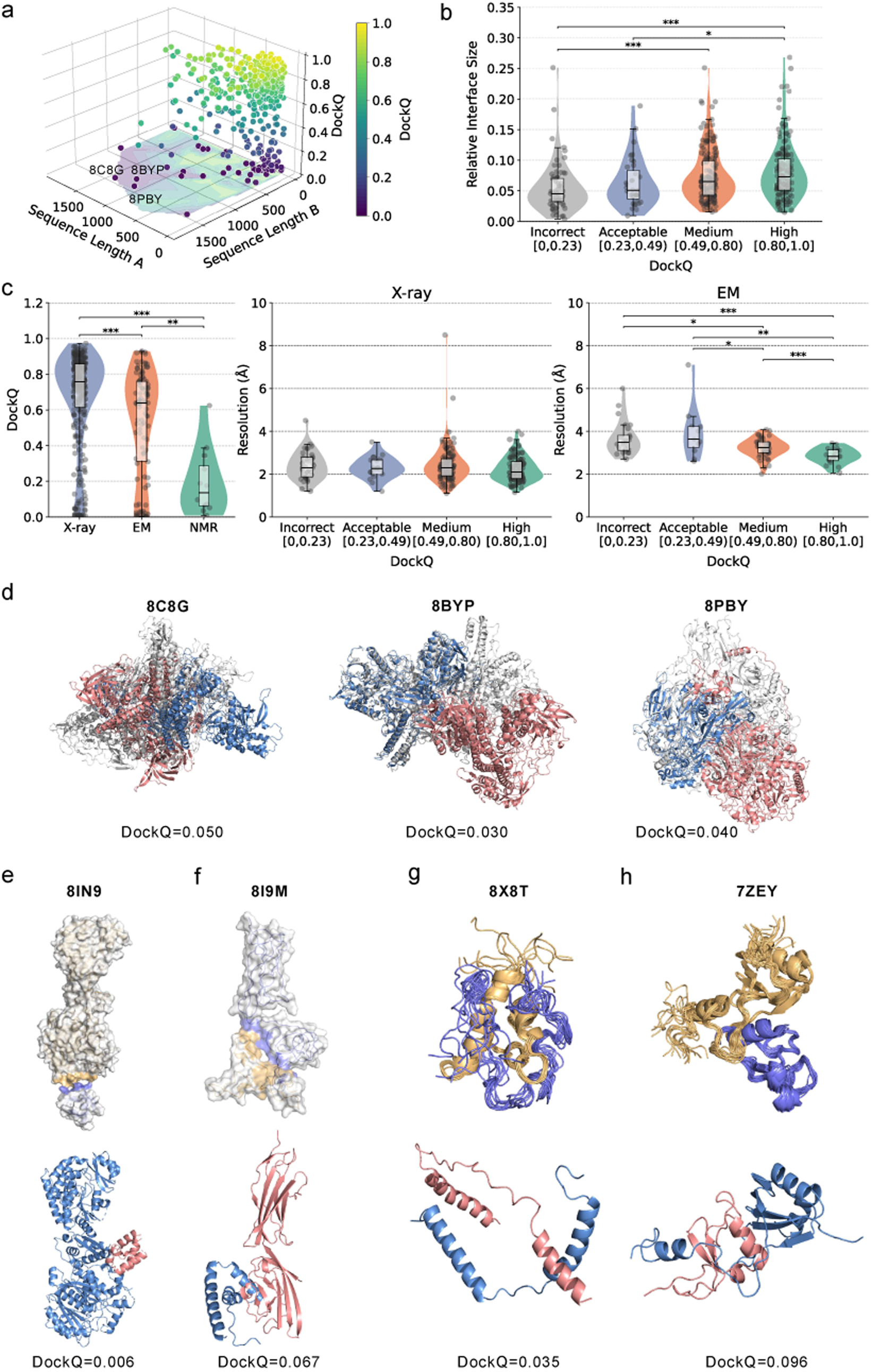
Analysis of factors influencing AF3 modeling accuracy and characterization of failure cases. (**a**) Three-dimensional scatter plot showing the relationship between the sequence lengths of the two chains and the resulting DockQ scores. (**b**) Distributions of relative interface sizes across four DockQ quality bins (Incorrect: < 0.23, Acceptable: 0.23∼0.49, Medium: 0.49∼0.8, and High: > 0.8). Relative interface size is defined as the ratio of the interface solvent accessible surface area (SASA) to the total SASA. (**c**) c (left), and resolution distributions across DockQ quality bins for X-ray and EM structures (center and right). Statistical significance (two-sided Wilcoxon rank-sum test): *: *P* < 0.05, **: *P* < 0.01, ***: *P* < 0.001. (**d**) Representative modeling failures for large complexes (total sequence length > 2,400 residues; > 1,000 residues per chain; PDB ID: 8C8G, 8BYP, and 8PBY). Experimental structures are gray; predictions are colored by chain. (**e-f**) Surface representations of experimental structures (top) and structures of predicted top-ranked models (bottom; Chain A: blue, Chain B: salmon) for 8IN9 (e) and 8I9M (f), with interface regions highlighted in slate and orange. (**g-h**) NMR ensembles (top) and structures of predicted top-ranked models (bottom; Chain A: blue, Chain B: salmon) for 8X8T (g) and 7ZEY (h).

#### Relative interface size

We defined relative interface size as the ratio of interface area to total solvent-accessible surface area and observed a certain correlation with prediction quality (Fig. 5b).

Complexes classified as incorrect or acceptable (DockQ < 0.49) exhibit smaller relative interfaces than those achieving medium or high accuracy. Small interfaces often reflect transient or weak interactions and/or enrichment in intrinsically disordered regions (IDRs). For example, the *Streptomyces graminofaciens* heterodimer 8IN9 features a highly transient interface involving fewer than 10 residues (Fig. 5e). While AF3 accurately folds the monomers (TM-score > 0.7), it fails to recover the docking pose (DockQ < 0.01). Similarly, the target 8I9M—dominated by IDRs and a minimal interface (3.6% of total area; Fig. 5f)—shows failures at both monomeric and quaternary levels. These cases indicate that limited surface complementarity, coupled with transient binding or disorder, severely hampers AF3’s interface inference.

#### Experimental structure determination method

We further assessed the influence of experimental structure determination methods on prediction accuracy (Fig. 5c). Statistically significant differences were observed in DockQ scores among structures solved by X-ray crystallography, cryo-electron microscopy (cryo-EM), and nuclear magnetic resonance (NMR). Notably, X-ray crystal structures yielded the highest overall DockQ scores, with accuracy being largely independent of experimental resolution. We attribute this robustness to two factors: first, the X-ray dataset is predominantly composed of high-resolution structures (mostly 1–3 Å) with well-defined interfaces, providing an essential benchmark for evaluating prediction accuracy [22]; and second, the overrepresentation of X-ray data in the AF3 training set likely biases the model’s priors toward the packing geometries and rigid conformations characteristic of crystal lattices.

In contrast, for cryo-EM targets, prediction quality correlates strongly with experimental resolution, where high-quality models are significantly associated with higher resolution inputs. This trend reflects the inherent characteristics of cryo-EM maps, where interface regions often exhibit lower local resolution compared to the protein core [23], thereby limiting prediction accuracy relative to X-ray targets. Finally, structures determined by NMR proved to be the most challenging, yielding the lowest median DockQ scores. This is likely because NMR structures often capture solution-state dynamics or contain flexible/disordered regions, which are intrinsically difficult for static inference models to resolve accurately (e.g., 8X8T, 7ZEY in Fig. 5g-h).

In conclusion, we identified that modeling failures are predominantly associated with large assemblies, restricted interface areas, and structures determined at low resolution. To enhance robustness, future training strategies should prioritize upsampling such challenging targets—including large assemblies, small-interface complexes, IDR-enriched interactions, and diverse experimental modalities—thereby addressing biases in current datasets and improving generalization for biologically relevant protein–protein interactions.

### Impact of MSA pairing to AlphaFold-Multimer

Benchmarking on the HD439 dataset revealed that AF3 consistently outperforms AlphaFold-Multimer (AFM) across all evaluated MSA configurations (Fig. 6a). Specifically, when employing *p*MSAs, AF3 achieved an average DockQ score of 0.613, representing a substantial margin over AFM (0.568). This performance gap is further illustrated in Supplementary Fig. 5a, where AF3 demonstrates superior accuracy across a broader spectrum of targets. These results are highly consistent with the benchmarks established in the AF3 paper.

**Fig. 6.**
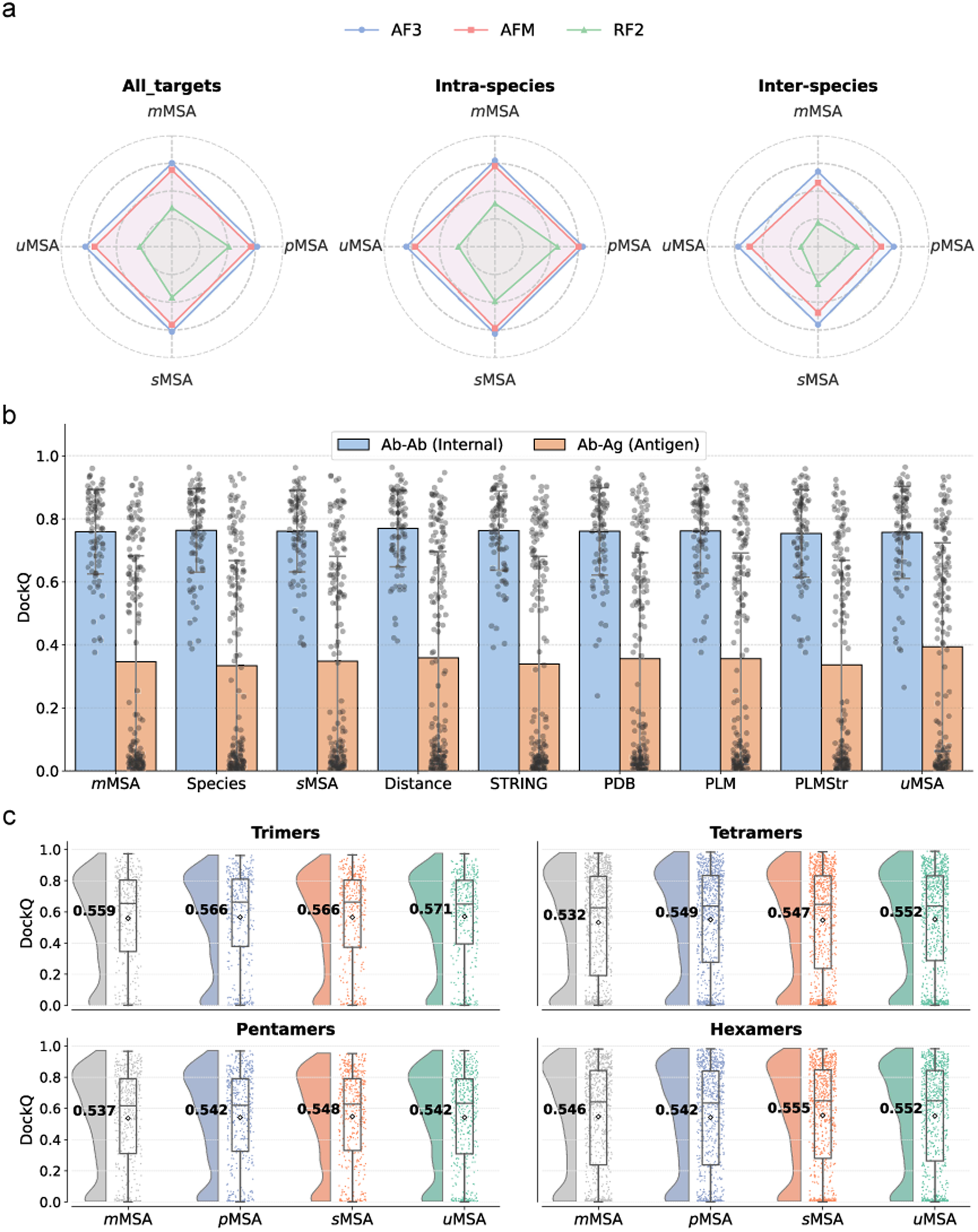
Benchmarking MSA strategies across different predictive models and complex types. (**a**) Radar charts comparing mean DockQ scores of AlphaFold3 (AF3), AlphaFold-Multimer (AFM), and RoseTTAFold2 (RF2) on HD439 and its subsets. Vertices represent the four MSA strategies. Detailed values are summarized in Supplementary Table 2. (**b**) Interface DockQ score distributions for 89 antibody-antigen complexes predicted by AF3 across various MSA strategies. Interfaces are categorized as internal antibody (Ab-Ab) or antibody-antigen (Ab-Ag). Bar heights represent mean values. Detailed values are summarized in Supplementary Table 3. (**c**) Interface DockQ score distributions for higher-order complexes (trimers, tetramers, pentamers, and hexamers) predicted by AF3. Diamonds and bold numerals indicate mean DockQ scores.

Regarding the impact of MSA pairing, AFM exhibits a trend similar to AF3: *s*MSA remains competitive with *p*MSA overall (0.562 vs. 0.568) and even surpasses *p*MSA on inter-species complexes (0.475 vs. 0.454). However, a notable difference arises in intra-species complexes, where the performance decline of *s*MSA relative to *p*MSA is more significant in AFM (0.587 vs. 0.601) compared to AF3 (0.627 vs. 0.632). Furthermore, unlike AF3, incorporating deep unpaired MSAs (*u*MSA) yields suboptimal results for AFM in intra-species cases.

In essence, MSA pairing appears more critical for AFM than for AF3. This disparity can be attributed to the varying degrees of reliance on MSA information inherent in their architectures. AFM maintains an explicit MSA representation throughout its entire 48-block Evoformer, employing extensive axial attention modules for updates. In contrast, AF3 processes MSA information using a streamlined MSA Module comprising only 4 blocks, devoid of axial attention operations. Consequently, AFM is significantly more sensitive to explicit pairing relationships in the input than AF3. This observation further substantiates our earlier analysis that sensitivity to MSA pairing is intrinsically linked to network architecture.

Moreover, we found that although the efficacy of *u*MSA is more variable for AFM, it exhibits distinct complementarity with *p*MSA (Supplementary Fig. 5b). Capitalizing on these divergent performance profiles, we evaluated a composite strategy that pools candidate models from *m*MSA, *p*MSA, and *u*MSA modes, selecting the top-ranked model based on confidence scores. This approach yielded a superior mean DockQ of 0.632 on the HD439 dataset. Notably, it boosted accuracy on the intra-species and inter-species subsets to 0.656 and 0.551, respectively, representing gains of 9.2% and 12.2% over the best-performing individual strategy for each category.

### Impact of MSA pairing to RoseTTAFold2

We extended our evaluation to another representative method, RoseTTAFold2 (RF2), using the HD439 benchmark (Fig. 6a). Compared to AF3 and AFM, RF2 exhibits lower overall accuracy and a more pronounced dependency on MSA pairing, with *p*MSA consistently outperforming *s*MSA and *u*MSA for both intra- and inter-species targets. This behavior can be attributed to the shallower architecture of RF2, which utilizes only 36 blocks for representation updates, in contrast to the ≥48 blocks employed by AF3 and AFM. This hypothesis is supported by our earlier analysis of AF3 (Supplementary Fig. 3), which demonstrated that inter-chain prediction accuracy continues to improve significantly beyond the 36th block.

These observations further confirm that the capacity to bypass explicit MSA pairing is intrinsically linked to neural network architecture and depth. Nevertheless, the *s*MSA strategy still substantially outperforms the *m*MSA baseline for both intra- and inter-species complexes, illustrating that even the relatively lightweight RF2 architecture retains the capability to recover inter-chain interactions from a pre-filtered pool of homologous sequences without strict pairing constraints.

### MSA pairing for antibody-antigen complexes

Antibody-antigen complexes are critical for immunological research and vaccine design but pose significant modeling challenges due to the hypervariability of CDR loops (or the lack of co-evolutionary constraints). We evaluated the impact of various MSA pairing and augmentation strategies on AF3 performance using a benchmark dataset of 89 antibody-antigen complexes derived from the DeepSCFold study.

As shown in Fig. 6b, the primary bottleneck lies in predicting the antibody-antigen interface. While all strategies successfully captured the interactions between antibody heavy and light chains (Ab-Ab; mean DockQ ∼0.8), accuracy for antibody-antigen (Ab-Ag) interfaces remained significantly lower, averaging between 0.3 and 0.4. Consequently, our subsequent analysis focuses exclusively on the antibody-antigen interfaces.

Compared to the *m*MSA baseline (DockQ 0.346), incorporating various pairing strategies yielded only marginal improvements (DockQ 0.334–0.359) for Ab-Ag interfaces (Supplementary Table 3); notably, some pairing methods (e.g., Species, STRING, and PLMStr) even proved detrimental to performance. This indicates that standard MSA pairing techniques offer limited utility for modeling antibody-antigen interactions. This finding underscores the distinct evolutionary nature of these complexes, where binding is driven primarily by somatic hypermutation rather than co- evolution. Furthermore, inter-species complexes constitute 74% of this dataset, a proportion significantly higher than in the HD439 set (23%). This compositional bias rationalizes the observation that *s*MSA (0.348) outperforms species-based *p*MSA (0.334).

Finally, augmenting the sequence pool via the UniProt database (*u*MSA) yielded the highest accuracy (DockQ: 0.394), surpassing the standard species-paired approach by 18.0% and the second-best method (Distance) by 9.7%. These results confirm that our general conclusion, that maximizing sequence depth is more effective than MSA pairing, extends to antibody-antigen complexes.

### MSA pairing for higher-order assemblies

We further extended our evaluation to a dataset of 491 higher-order hetero-oligomers, ranging from trimers to hexamers (Fig. 6c). The observations are largely consistent with the dimer-based benchmarks. Specifically, *p*MSA offered only marginal gains over *m*MSA, and shuffling the pairing (*s*MSA) did not compromise AF3 performance. Notably, *u*MSA remained a highly effective strategy, yielding DockQ scores that matched or outperformed *p*MSA across diverse oligomeric states.

Stratification into intra- and inter-species complexes further confirmed that inter-species targets remain significantly more challenging (Supplementary Fig. 5c). For intra-species trimers through pentamers, *u*MSA yielded average performance comparable to *p*MSA and *s*MSA. For inter-species trimers to pentamers, increasing MSA depth via unpaired homologs provided a slight performance gain. An apparent anomaly was observed in inter-species hexamers, where *p*MSA (DockQ = 0.472) appeared marginally superior to *u*MSA (0.469); however, this discrepancy is attributed to outliers corresponding to two interfaces of the target with PDB ID of 8AIL (Supplementary Fig. 5d). In fact, AF3 using *u*MSA successfully sampled high-quality structures (best model DockQ = 0.905), yet these were not selected as the top-ranking models (Supplementary Fig. 5e). Collectively, our results indicate that augmenting MSA depth is also an effective strategy for the robust prediction of higher-order hetero-oligomers, though challenges in effective model selection for the challenging targets persist.

## CONCLUSIONS

Protein-protein complex structure prediction has long been a formidable challenge. To address this, substantial effort has been dedicated to optimizing MSA pairing to extract inter-chain co-evolutionary signals, a step long regarded as indispensable for accurate prediction. However, the advent of AlphaFold models, particularly AF3, has fundamentally altered this paradigm.

Our systematic benchmarking demonstrates that for AF3, disrupting explicit pairing patterns does not degrade performance; in fact, it enhances accuracy for inter-species complexes. This finding challenges the prevailing dogma that strict phylogenetic pairing is essential for complex prediction. We attribute this robustness to the intrinsic geometric and physicochemical complementarity between subunits, as well as the AF3 network architecture, which can autonomously recover interaction patterns.

Instead, we propose that the critical determinant is actually MSA depth. We demonstrated that by utilizing deeper, unpaired MSAs (*u*MSA), AF3 effectively surpasses the performance achieved with elaborate pairing schemes. This ‘depth-over-pairing’ principle is particularly transformative for challenging scenarios, such as antibody-antigen interactions and inter-species complexes, where matching orthologs are naturally scarce. The validity of this conclusion is further substantiated by the recent success of the omicMSA strategy.

Furthermore, our study highlights a clear correlation between modeling difficulties and factors such as the complex size, interface sizes, and experimental resolution. In light of these findings, we propose that the critical path for future improvement involves prioritizing the depth and quality of monomeric MSAs and adopting ‘up-sampling’ protocols to better address these underrepresented targets.

## METHODS

### 1. Construction of benchmark datasets

#### Initial data curation

We collected a dataset of heteromeric protein complexes (2–6 chains; ≥30 residues per chain) released after the training set date for AF3 and AFM (September 30, 2021), sourced from the protein–protein interaction dataset in Q-BioLiP [24]. To ensure structural quality and biological relevance, we applied the following inclusion criteria: (1) exclusion of complexes containing peptides or nucleic acids (DNA/RNA); (2) resolution better than 10 Å; (3) absence of chains with unknown residues; (4) a requirement for a connected interaction graph, defined by the presence of at least one inter-chain atomic contact ≤8 Å for every chain; (5) a maximum total sequence length of 4,000 residues, with individual chains limited to 1,500 residues; and (6) exclusion of antibodies and *de novo* designed proteins. This filtering process yielded 6,832 candidate complexes, comprising 2,771 dimers, 1,266 trimers, 1,249 tetramers, 886 pentamers, and 660 hexamers.

#### Exclusion of training set redundancy

To prevent data leakage from the AF3 and AFM training set (PDB structures released before September 30, 2021), we implemented a two-step clustering protocol [4]. First, chains from both the benchmark and training datasets were pooled and clustered at a 70% sequence identity threshold using CD-HIT [25]. Second, two interfaces were considered redundant if their constituent chain clusters were identical (invariant to ordering). We removed any complex from our dataset containing an interface that mapped to the same interface cluster as any structure in the AlphaFold training set. This step reduced the dataset to 875 complexes.

#### Internal redundancy reduction

Finally, to eliminate redundancy within the benchmark itself, we performed hierarchical clustering separately for each oligomeric state. Using the interface-clustering logic described above, complexes with identical interface clusters were grouped, and the structure with the highest resolution was selected as the representative. The final benchmark dataset consists of 930 heteromeric complexes, including 439 dimers, 145 trimers, 187 tetramers,70 pentamers, and 89 hexamers.

### 2. Monomer MSA generation

Candidate MSAs for individual chains were generated using two distinct protocols. (1) HHblits-based generation: Searches were performed against the Uniclust30 (2018_08) database using HHblits [26]. We generated up to 12 MSAs from this search with various cutoffs for E-value (10^−40^,10^−10^,10^−3^, 1), coverage (50%, 75%), and identity (90%, 100%). (2) MMseqs2-based generation: Searches were conducted against the UniRef30 (2022_02) and ColabFold environmental (2021_08) databases using MMseqs2 [27, 28]. The resulting alignments were merged and filtered using HHfilter with a 99% sequence identity threshold. For each chain in hetero-multimeric complexes, the optimal MSA was identified based on the maximal mean pLDDT score from monomeric AF2 predictions. Subunit MSAs were subsequently concatenated in a dense format for AF3 or a block-diagonal format for AFM. The final resulting alignments are denoted as the *m*MSA.

### 3. MSA pairing methods

To ensure consistent comparisons, all paired MSAs were derived from identical monomeric alignments generated for each subunit using JackHMMER [29] searches against the UniProt database (version 2020_05), retaining a maximum of 50,000 hits. We evaluated six strategies for concatenating subunit alignments to form paired MSAs for hetero-multimers:

- **Species** [11, 21]: concatenating sequences from the same species using the ‘top-down’ approach consistent with AFM;
- **Distance** [21, 30]: pairing sequences with UniProt accession indices differing by fewer than 10;
- **STRING** [21, 31]: pairing sequences with interaction confidence scores >0.5 in the STRING database;
- **PDB** [21, 32]: pairing sequences mapped to the same PDB entry;
- **PLM** and **PLMStr**: employing the DeepSCFold framework [15], which ranks monomeric hits by sequence (PLM) or structural (PLMStr) similarity, followed by subunit concatenation based on interaction probabilities predicted via a deep neural network.

Implementation of the non-species-based strategies followed the DeepSCFold codebase. For AF3 predictions, the generated paired MSAs were appended to the unpaired monomeric MSAs (*m*MSA) to constitute the final input.

### 4. Strategies for MSA shuffling and sequence augmentation

To evaluate the efficacy of the species-based MSA pairing strategy (default pairing) in predicting heteromeric complex structures, we constructed a control alignment termed “*s*MSA”. Specifically, we randomly shuffled the sequence order within each chain’s original paired MSA (excluding the query sequence), concatenating them laterally and merging them with the monomeric MSAs. This procedure disrupts specific inter-chain pairing patterns while retaining the identical monomeric sequence content of the original paired MSA.

Furthermore, to maximize the utility of sequences derived from UniProt MSAs without pairing, we generated two new MSAs using the following protocols: (1) merging the UniProt MSA with the *m*MSA, followed by filtering using HHfilter with a sequence identity threshold of 99%; and (2) filtering the UniProt MSA to exclude sequences sharing >90% or <25% identity with the query, as well as removing sequences identical to those in the *m*MSA, then merging with *m*MSA. These two sequence-augmented MSAs (denoted as *u*MSAs) served as distinct inputs for subsequent structure prediction.

### 5. Configurations for running AF3, AFM, and RF2

**AlphaFold3 (AF3).** Structure predictions were performed using the official implementation of AF3 [4] without structural templates. For the HD439 dataset, 75 predicted structures were generated for each input MSA using 15 distinct random seeds (five models per seed). To ensure comparability, the same set of 15 seeds was maintained across all MSA variants for a given complex. For antibody-antigen complexes and higher-order multimers, five distinct random seeds were used to accelerate sampling.

**AlphaFold-Multimer (AFM).** The AFM results are generated using AlphaFold v2.3.2 [2], modifying the standard inference script to input the four pre-constructed MSAs into the network. During modeling, relaxation was turned off, as it has been verified that final relaxation does not improve model accuracy [33]. No templates were used, and five predictions were generated from five multimer models (one seed). Additionally, we set the hyperparameter random seed to a fixed value to ensure that AFM is deterministic, producing the same prediction when given the same MSA input.

**RoseTTAFold2 (RF2).** RF2 [3] was obtained from the official repository (https://github.com/uw-ipd/RoseTTAFold2). For heterodimers in the HD439 dataset, structure predictions were generated by feeding pre-constructed complex MSAs directly into the inference script (network/predict.py). To ensure deterministic output, network randomness was fixed, while all other parameters were maintained at their default settings.

### 6. DockQ for measuring the accuracy of complex models

To evaluate the interface quality of protein complexes modeled by AF3, AFM, and RF2, we adopted the DockQ v2.0 [34]. All DockQ metrics were calculated using the official implementation available at https://github.com/bjornwallner/DockQ/.

## DATA AVAILABILITY

All data used in this work, including protein sequences, native protein structures, input MSAs, and predicted structure models are available at: https://yanglab.qd.sdu.edu.cn/MSApair/.

## CODE AVAILABILITY

Codes for running AlphaFold3, AlphaFold-Multimer, and RoseTTAFold2 were downloaded from the respective URLs in GitHub.

## AUTHOR CONTRIBUTIONS

J.Y. conceived and supervised the project; Y.L. designed and implemented the experiments; W.W. analyzed the experimental results and conducted the theoretical analysis of the AF3 modeling. W.W. and Z.P. co-supervised the study; All authors reviewed and approved the final version of the paper.

## ACKNOWLEDGEMENTS

This work was supported by the National Natural Science Foundation of China (NSFC T2225007, 32430063, 62501364), the National Key Research and Development Program of China (2024YFA0916901), the Postdoctoral Fellowship Program and the China Postdoctoral Science Foundation (BX20240212, 2025M783122), and the Fundamental Research Funds for the Central Universities.

