## Supplementary Material for "Benchmarking MSA pairing for protein-protein complex structure prediction reveals a depth-over-pairing principle"

### Supplementary Information

#### Supplementary Note 1: Mathematical intuition for inter-sequence information propagation in AF3

In this section, we provide a simplified mathematical analysis to elucidate how AF3 extracts latent co-evolutionary patterns from unpaired MSAs. We demonstrate that the network architecture naturally facilitates interactions between signals derived from different sequences (e.g., sequence *a* and sequence *b*), thereby bypassing the strict requirement for row-wise alignment. For tractability, auxiliary components (linear layers, LayerNorm, activations) are omitted.

##### 1. Assimilation of MSA signals via outer-product-mean

The initial transfer of information from the MSA representation (**m**) to the pair representation (**z**) occurs via the outer-product-mean operation. For a residue pair $(i, j)$, the network aggregates outer products (⊗) across all available sequences $s \in \{1, \dots, N_{seq}\}$:

$$\mathbf{z}_{ij} = \mathbf{z}_{ij} + \frac{1}{N_{seq}} \sum_s \mathrm{Linear}(\mathbf{m}_i^{(s)}) \otimes \mathrm{Linear}(\mathbf{m}_j^{(s)}) \quad (1)$$

Simplifying this, the pair representation $\mathbf{z}_{ij}$ effectively becomes a summation of intra-sequence correlations:

$$\mathbf{z}_{ij} \sim \mathbf{z}_{ij} + \frac{1}{N_{seq}} \sum_s \mathbf{m}_i^{(s)} \otimes \mathbf{m}_j^{(s)} = \mathbf{z}_{ij} + \frac{1}{N_{seq}} \left( \mathbf{m}_i^{(a)} \otimes \mathbf{m}_j^{(a)} + \mathbf{m}_i^{(b)} \otimes \mathbf{m}_j^{(b)} + \cdots \right) \quad (2)$$

##### 2. Cross-sequence mixing via triangle updates

The core "reasoning" capability of AF3 lies in the triangle multiplicative update and triangle attention modules. Both mechanisms update the edge $\mathbf{z}_{ij}$ by integrating information from a third node *k* involving a multiplicative interaction (e.g., $\mathbf{z}_{ik} \odot \mathbf{z}_{kj}$).

Substituting the aggregated form from Eq. (2) into this multiplicative update reveals the emergence of cross-sequence terms. Considering the interaction between edges $\mathbf{z}_{ik}$ and $\mathbf{z}_{kj}$:

$$\mathrm{Update}(\mathbf{z}_{ij}) \propto \mathbf{z}_{ik} \odot \mathbf{z}_{kj}$$

$$\propto \left[ \sum_s \mathbf{m}_i^{(s)} \otimes \mathbf{m}_k^{(s)} \right] \odot \left[ \sum_s \mathbf{m}_k^{(s)} \otimes \mathbf{m}_j^{(s)} \right] \quad (3)$$

Expanding this product yields two types of terms:

- Intra-sequence terms (where $a = b$): Traditional co-evolutionary signals.
- Inter-sequence "Cross" terms (where $a \neq b$):

$$\mathcal{T}_{\text{cross}} = \left(\mathbf{m}_i^{(a)} \otimes \mathbf{m}_k^{(a)}\right) \odot \left(\mathbf{m}_k^{(b)} \otimes \mathbf{m}_j^{(b)}\right) \quad (4)$$

Eqs. (3) and (4) demonstrate that through the triangle updates, the network mathematically forces an interaction between the evolutionary history of sequence $a$ (at positions $i, k$) and sequence $b$ (at positions $k, j$). This proves that AF3 is not limited to analyzing single rows of the MSA implicitly; rather, it synthesizes a global consistency landscape by mixing signals across the entire depth of the MSA. This explains our empirical finding that the AF3 achieves high accuracy independent of elaborate row-wise pairing.

#### Tables

**Supplementary Table 1. Comparison of DockQ scores for 14 protein-protein interactions predicted by AF3 across diverse MSA configurations.** The evaluated inputs include *m*MSA, *p*MSA, *s*MSA, and *o*MSA generated using the omicMSA pipeline, alongside a baseline *u*MSA derived from the UniProt database. *Values in parentheses denote reference DockQ scores of the best models available from the Human PPI benchmark (http://prodata.swmed.edu/humanPPI) generated using AF2/AFM.

| UniProt Accession | PDB ID | *m*MSA | *p*MSA* | *s*MSA | *o*MSA | *u*MSA |
|---|---|---|---|---|---|---|
| O14744_Q9BQA1 | 6UGH | 0.873 | 0.848(0.856) | 0.861 | 0.862 | 0.850 |
| P68371_Q71U36 | 8SH7 | 0.013 | 0.581(0.705) | 0.542 | 0.582 | 0.476 |
| Q13418_Q9NVD7 | 6MIB | 0.923 | 0.925(0.907) | 0.909 | 0.913 | 0.924 |
| P17535_P53539 | 7UCC | 0.659 | 0.644(0.563) | 0.624 | 0.657 | 0.673 |
| P23471_Q12860 | 3S97 | 0.119 | 0.861(0.795) | 0.848 | 0.814 | 0.853 |
| P62837_Q96EQ8 | 8GBQ | 0.854 | 0.865(0.847) | 0.885 | 0.885 | 0.496 |
| Q14213_Q8NEV9 | 8XWY | 0.039 | 0.763(0.752) | 0.815 | 0.747 | 0.744 |
| O14944_P00533 | 7LEN | 0.799 | 0.823(0.704) | 0.804 | 0.869 | 0.804 |
| O00258_O43681 | 8CQZ | 0.863 | 0.923(0.702) | 0.832 | 0.912 | 0.869 |
| P05556_P08648 | 4WK0 | 0.778 | 0.776(0.778) | 0.786 | 0.816 | 0.799 |
| P08195_Q9UPY5 | 7EPZ | 0.538 | 0.473(0.530) | 0.438 | 0.545 | 0.539 |
| Q15019_Q92599 | 6UPR | 0.883 | 0.897(0.885) | 0.004 | 0.900 | 0.004 |
| O43520_Q3MIR4 | 7VGH | 0.610 | 0.629(0.753) | 0.641 | 0.602 | 0.637 |
| Q13416_Q9UBD5 | 5UJ8 | 0.596 | 0.579(0.565) | 0.585 | 0.595 | 0.581 |
| Average | | 0.611 | 0.756(0.739) | 0.684 | 0.764 | 0.661 |

**Supplementary Table 2. DockQ scores of AlphaFold3 (AF3), AlphaFold-Multimer (AFM), and RoseTTAFold2 (RF2) on HD439 and its subsets.**

| | **Methods** | ***m*MSA** | ***p*MSA** | ***s*MSA** | ***u*MSA** |
|---|---|---|---|---|---|
| HD439 | AF3 | 0.602 | 0.613 | 0.612 | 0.623 |
| | AFM | 0.552 | 0.568 | 0.562 | 0.555 |
| | RF2 | 0.281 | 0.410 | 0.365 | 0.232 |
| Intra-species | AF3 | 0.620 | 0.632 | 0.627 | 0.638 |
| | AFM | 0.579 | 0.601 | 0.587 | 0.573 |
| | RF2 | 0.312 | 0.449 | 0.393 | 0.264 |
| Inter-species | AF3 | 0.540 | 0.545 | 0.561 | 0.574 |
| | AFM | 0.459 | 0.454 | 0.475 | 0.491 |
| | RF2 | 0.174 | 0.278 | 0.268 | 0.123 |

**Supplementary Table 3. Average interface DockQ scores of AlphaFold3 (AF3) on 89 antibody-antigen complexes.** Interfaces are categorized as internal antibody (Ab-Ab) or antibody-antigen (Ab-Ag). Bold values represent the best DockQ in each category.

| **Methods** | **Ab-Ab (Internal)** | **Ab-Ag (Antigen)** |
|---|---|---|
| *m*MSA | 0.759 | 0.346 |
| Species | 0.764 | 0.334 |
| *s*MSA | 0.761 | 0.348 |
| Distance | **0.770** | 0.359 |
| STRING | 0.763 | 0.339 |
| PDB | 0.760 | 0.356 |
| PLM | 0.762 | 0.356 |
| PLMStr | 0.754 | 0.337 |
| *u*MSA | 0.757 | **0.394** |

### Figures

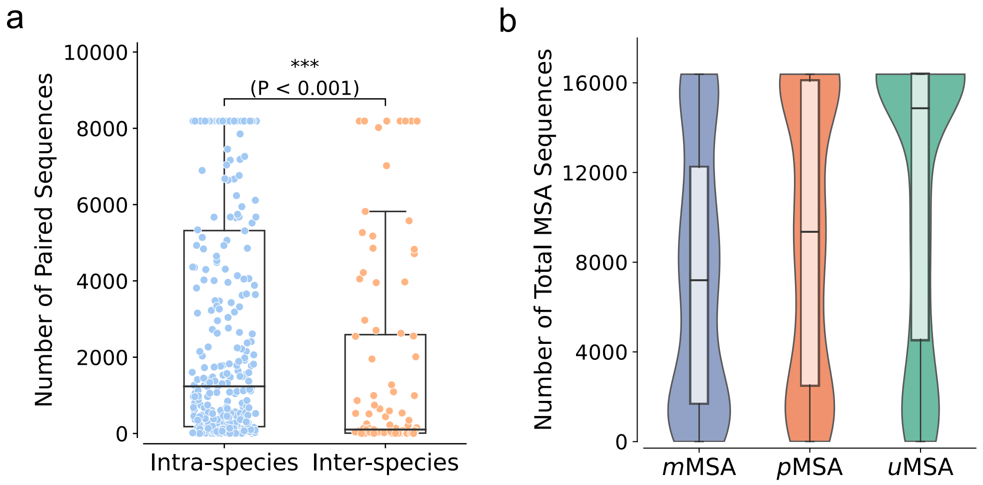


**Supplementary Fig. 1. Distribution of the sequence number of different MSAs on the HD439 dataset.** (**a**) Distribution of the number of paired sequences generated by *p*MSA (Species-based) for the intra- and inter-species subsets. (**b**) Distribution of the total number of sequences in *m*MSA, *p*MSA, and *u*MSA.

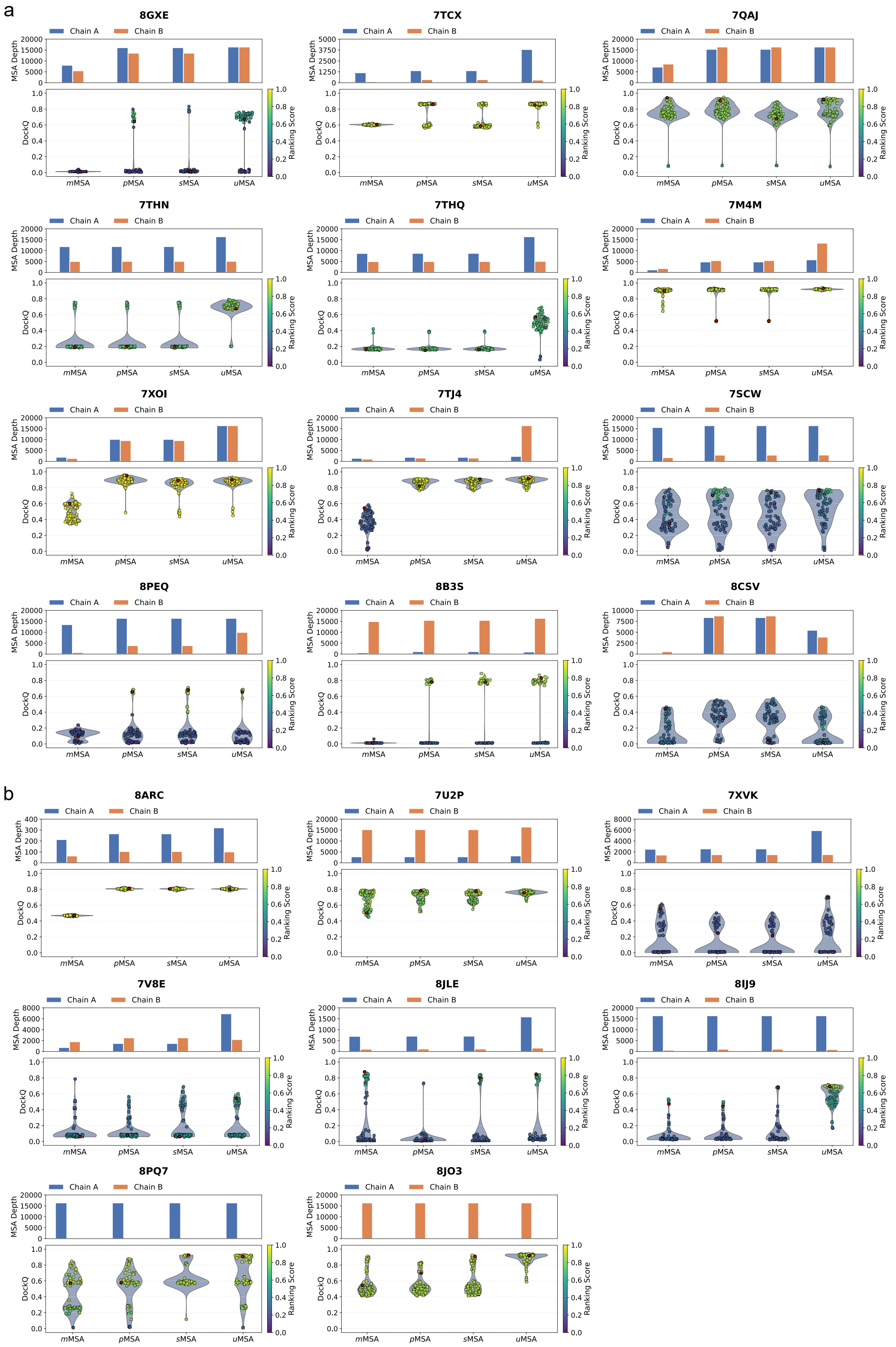


**Supplementary Fig. 2. Visualization analysis for the 20 examples exhibiting significant performance variations (DockQ gap > 0.2) across the four MSA strategies.** Top panels: Bar charts showing the effective MSA depth (number of sequences) for each chain. Bottom panels:

Violin plots displaying the distribution of DockQ scores for 75 generated models. Red dots indicate the DockQ scores of the top-ranked models selected by the confidence metric. (**a**) Intra-species dimers. (**b**) Inter-species dimers.

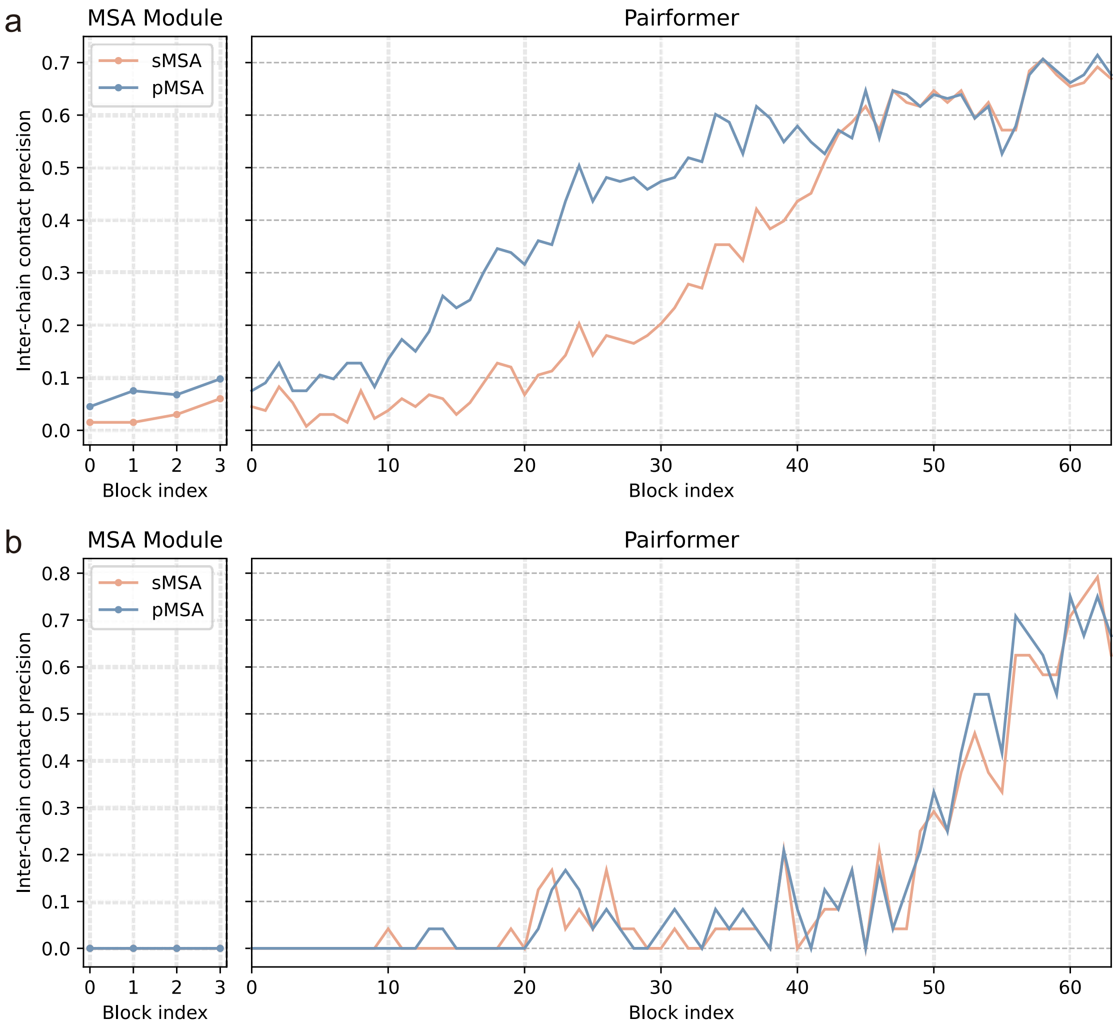


**Supplementary Fig. 3. Probing the progression of inter-chain contact information through AF3 blocks.** This analysis quantifies the emergence of contact signals across network blocks using 68 logistic regression models, which map the output pair representations from each block to inter-residue contact probabilities. These models were trained on 100 protein monomers randomly selected from the trRosetta training set [1] (released prior to May 2018). Performance is evaluated based on the precision of the top-10 predicted inter-chain contacts. Given the licensing restrictions associated with the official AF3 model parameters, we conducted this analysis using Boltz-2 [2], an open-source PyTorch implementation of the AF3 architecture. (**a**) An intra-species complex (PDB ID: 7PU4). (**b**) An inter-species complex (PDB ID: 7B57).

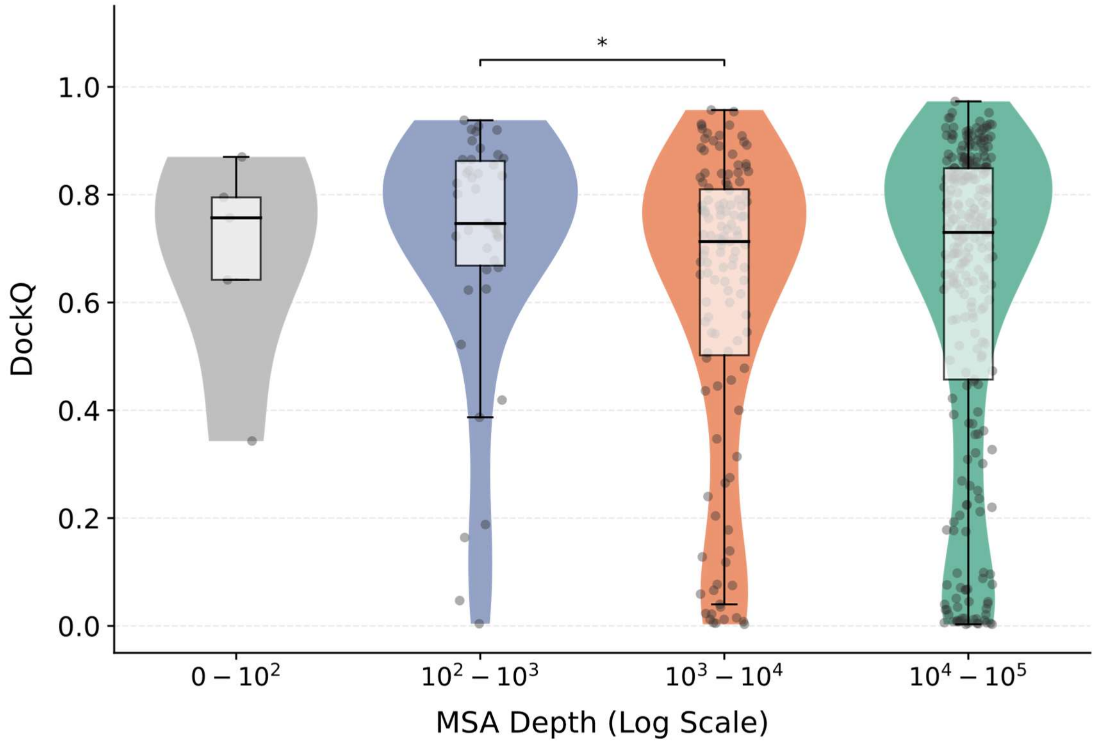


**Supplementary Fig. 4. Distribution of DockQ scores for predicted models, binned by MSA depth.** Statistical significance was assessed using a two-sided Wilcoxon rank-sum test (*: $P < 0.05$, **: $P < 0.01$, ***: $P < 0.001$)

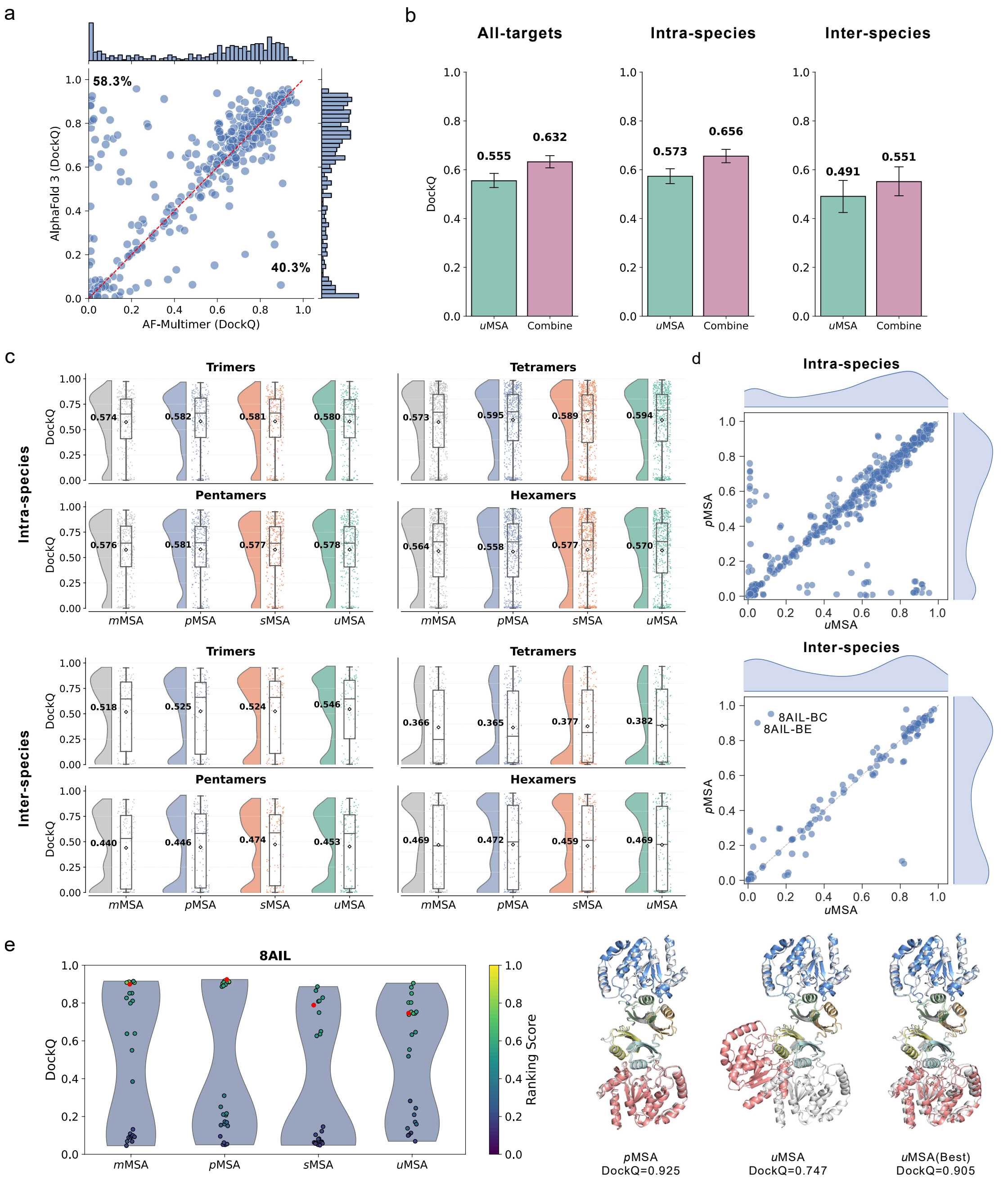


**Supplementary Fig. 5. Benchmarking MSA strategies across different predictive models and complex types.** (**a**) Head-to-head comparison of AlphaFold 3 and AlphaFold-Multimer using *p*MSA input on the HD439 dataset. The percentage represents the proportion of targets for which the corresponding method is superior (DockQ gap > 0). (**b**) Performance comparisons of *u*MSA and Combine (a composite strategy that pools candidate models from *m*MSA, *p*MSA, and *u*MSA modes, selecting the top-ranked model based on confidence scores) using AFM on the HD439 dataset. (**c**) Interface DockQ score distributions on intra- and inter-species complexes for higher-order complexes (trimers, tetramers, pentamers, and hexamers) predicted by AF3. Diamonds and bold numerals indicate mean DockQ scores. (**d**) Head-to-head comparisons of *u*MSA and *p*MSA for hexamers on intra- and inter-species complexes. (**e**) DockQ score distributions of 75 models and predicted structures across various MSAs for 8AIL. Experimental structures are shown in gray cartoon, while predicted structures are colored by chain.
